# *Bacteroides thetaiotaomicron*-infecting bacteriophage isolates inform sequence-based host range predictions

**DOI:** 10.1101/2020.03.04.977157

**Authors:** Andrew J. Hryckowian, Bryan D. Merrill, Nathan T. Porter, William Van Treuren, Eric J. Nelson, Rebecca A. Garlena, Daniel A. Russell, Eric C. Martens, Justin L. Sonnenburg

## Abstract

Our emerging view of the gut microbiome largely focuses on bacteria and less is known about other microbial components such as of bacteriophages (phages). Though phages are abundant in the gut, very few phages have been isolated from this ecosystem. Here, we report the genomes of 27 phages from the United States and Bangladesh that infect the prevalent human gut bacterium *Bacteroides thetaiotaomicron*. These phages are mostly distinct from previously sequenced phages with the exception of two, which are crAss-like phages. We compare these isolates to existing human gut metagenomes, revealing similarities to previously inferred phages and additional unexplored phage diversity. Finally, we use host tropisms of these phages to identify alleles of phage structural genes associated with infectivity. This work provides a detailed view of the gut’s “viral dark matter” and a framework for future efforts to further integrate isolation- and sequencing-focused efforts to understand gut-resident phages.

## Introduction

Bacteriophages (phages) are highly abundant constituents of free-living and host-associated microbial communities (microbiomes) (Brussow and Hendrix, 2002, Barr et al., 2013). Like other microbiome members (e.g. bacteria, fungi), the diversity and abundance of phages differ between healthy and diseased individuals. While some gut resident phages appear to be unique to individual humans and stable across long time scales (Shkoporov et al., 2019), others correlate with host disease status (Manrique et al., 2016, Duerkop et al., 2018). These observations highlight the possibility that phages of host-associated microbiomes play central roles in the structure and function of these communities and may therefore impact human health. Taken together with the burgeoning antibiotic resistance crisis, this possibility amplifies the importance of phage therapy as an alternative or supplement to existing paradigms of microbiome management (e.g. widespread antibiotic use).

In contrast to the enthusiasm for phages and phage-based therapeutics is an underlying reality that gut-resident phages, as a microcosm of the global phage population, are poorly understood. This deficiency is highlighted when a typical path for generating and addressing hypotheses in the microbiome field is considered. For example, sequencing-based approaches are often used in the microbiome field to generate hypotheses about complex microbial ecosystems. These hypotheses are subsequently addressed experimentally using a varied and expanding toolkit of molecular, genetic, immunological, and microbiological tools. This sequencing-to-mechanism approach is not straightforward for phages for several reasons. First, unlike bacteria, phages do not have conserved marker genes (e.g. the 16S rRNA marker gene) that enable phylogenetic classification and analysis. Instead, phage genomes must be inferred from metagenomic studies, either based on conservation of phage-like genes (e.g., terminase, DNA polymerase) (Grazziotin et al., 2017), sequence identity relative to known phage isolates (Roux et al., 2015), or by database-independent approaches (Ren et al., 2017). While powerful for general characterization of changes in the composition of phage communities, inter-study methodological variation (e.g. sample preparation, contig assembly, reference databases used) can impact a study’s conclusions to a greater extent than the treatment effects (e.g. health or disease status) (Gregory et al., 2019).

Furthermore, metagenomic approaches fail to provide definitive information on the bacterial hosts of these phages. To address this deficiency, many methods have been developed to predict the bacterial hosts of phages inferred from metagenomes. For example, homology searches, identification of CRISPR spacers, and co-occurrence analysis were used to make the prediction that the highly prevalent and abundant crAssphage infects bacteria in the phylum *Bacteroidetes* (Dutilh et al., 2014). This prediction was validated in part when a crAss-like phage (CrAss001) was isolated on *Bacteroides intestinalis* (Shkoporov et al., 2018). However, based on the divergence of CrAss001 from the prototypical crAssphage combined with the astounding diversity of crAss-like phages across host associated and environmental microbiomes (Yutin et al., 2018, Guerin et al., 2018), it is likely that other crAss-like phages infect other bacterial strains within the *Bacteroidetes* phylum. Furthermore, crAss-like phages can simultaneously be biomarkers of healthy and diseased states. For example, one crAss-like phage, IAS virus, is enriched in HIV^+^ individuals with low CD4 counts (Oude Munnink et al., 2014) while some crAss-like phages are stable over a 12 month period in healthy humans (Shkoporov et al., 2019).

CrAss001 is one of four isolated and sequenced phages confirmed to infect *Bacteroides*, the most abundant bacterial genus in the human gut microbiome. The other three phages are B40-8 and B124-14 (which infect *B. fragilis)* and Hankyphage, which is present as a prophage in many *Bacteroides* strains (Benler et al., 2018, Ogilvie et al., 2012, Hawkins et al., 2008). Despite the prevalence of Hankyphage lysogens in the public databases, Hankyphage induced from a Hankyphage-containing *B. dorei* lysogen was unable to form plaques on Hankyphage-naïve *B. dorei* or additional *Bacteroides* species (Benler et al., 2018). Additionally, CrAss001 does not form robust plaques on *B. intestinalis* despite persisting at high levels in co-culture with its host for nearly one month (Shkoporov et al., 2018). These observations suggest that unexplored factors influence *Bacteroides-phage* interactions.

Additional *Bacteroides* phages were previously isolated but not sequenced, representing limitations in connecting experiment-focused studies with bioinformatics-focused studies. Our recent work incorporated 71 phage isolates to show that multiple phase-variable mechanisms, including capsular polysaccharides (CPS), modify bacteriophage susceptibility in *B. thetaiotaomicron*. These findings mirror work in other isolated phages where cps is a major determinant of phage-host tropism (Porter et al., 2019). However, in the absence of genome sequences, phage-encoded determinants of host tropism were not previously explored.

Here, we report the genomes of 27 phages that *infect B. thetaiotaomicron* (18 previously described isolates (Porter et al., 2019) and 9 new isolates; **Table S1**). By comparing these genomes with those of existing *Bacteroides* phage isolates and with phage genomes identified from publicly available metagenomic studies, we simultaneously reveal similarities to previously inferred phage genomes and additional unexplored phage diversity. Finally, analysis of these genomes in the context of their cps-specific host ranges reveals targets for future study aimed at understanding the structure-function relationship of phage host range and phylogeny. We suggest the utility and feasibility of future efforts that integrate both isolation- and computational-based methods. Such an approach would enrich databases of known phage-host pairs by providing additional reference genomes and definitive host information. Furthermore, isolated phages will enable investigators to build experimental systems to test hypotheses and theoretical predictions and contribute to a growing collection of phages that may be used in the future for therapeutic or biotechnological applications.

## Results

### Isolation and comparative analysis of 27 phages infecting B. thetaiotaomicron

Our study centers of phages isolated from four geographic locations, three within the US and one in Bangladesh (**Figs. 1A, Table S1**). Using a previously reported protocol for phage isolation (Porter et al., 2019), we isolated 9 bacteriophages from primary wastewater effluent from the Sand Island Wastewater Treatment Plant (Honolulu, Hawaii) or from sewer-adjacent pond water at two locations in Dhaka, Bangladesh. High titer stocks were prepared of these 9 phage isolates and of a subset of 17 phages from an existing collection of 71 *B. thetaiotaomicron-infecting* phages isolated from Ann Arbor, Michigan and San Jose, California (Porter et al., 2019). Phage genomes were sequenced and assembled. The phages are grouped into three genomically related clusters (α, β, γ) and have genomes that are on average 38kb +/- 0.4kb, 99kb +/- 0.3kb, and 177kb +/- 4.5kb, respectively, and exhibit extensive genomic mosaicism (**Figs. S1-S3; Table S1**). Transmission electron microscopy of one representative from each cluster reveals distinct virion morphologies. Based on these representatives, cluster α phages are siphoviruses, cluster β phages are podoviruses, cluster γ phages are myoviruses, and the capsid sizes of these phages scale with genome size (**Figure 1B-D**).

**Figure 1.**
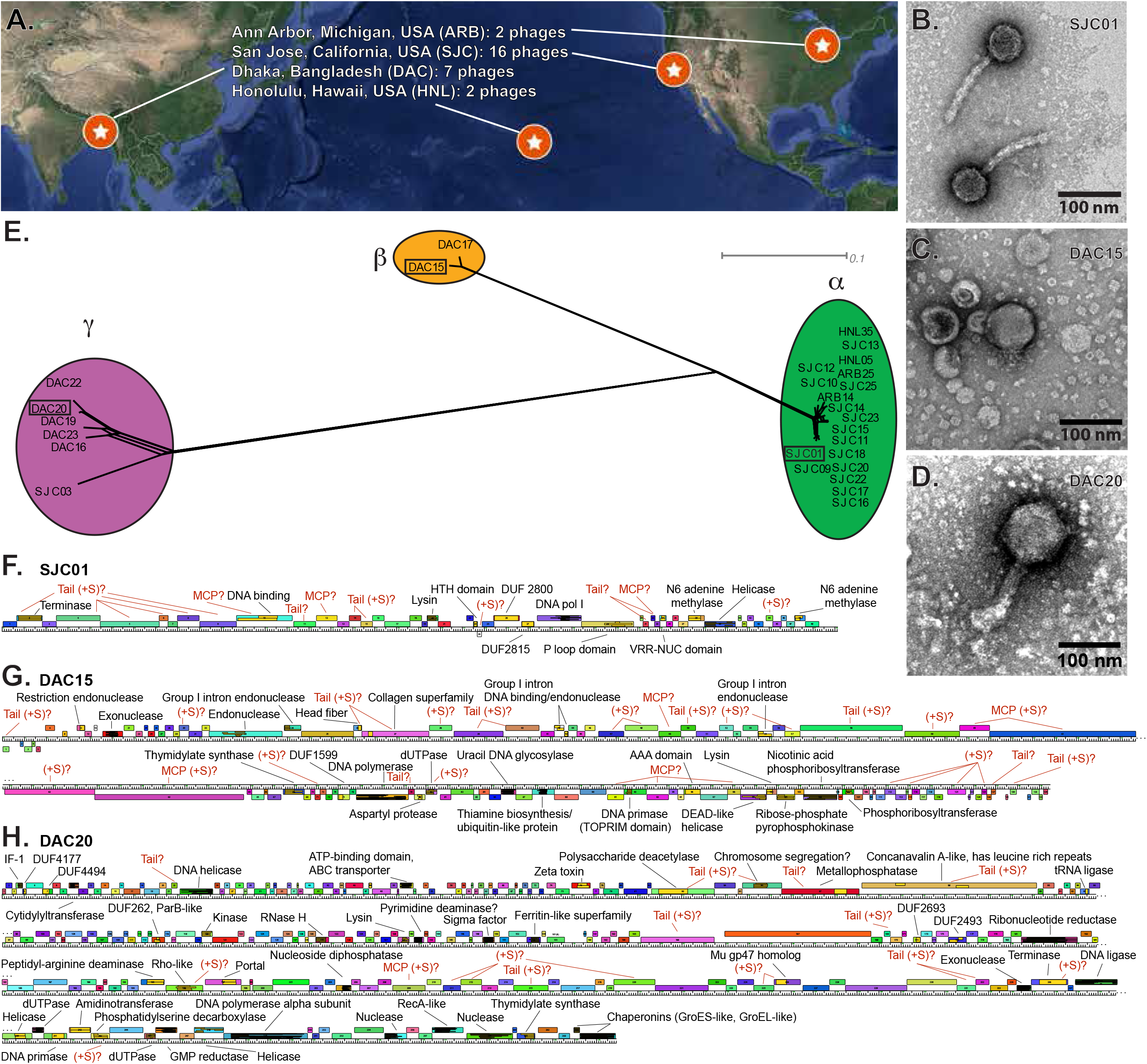
Isolation and characterization of 27 *Bacteroides thetaiotaomicron*-infecting phages. (**A**) Phages were isolated from wastewater samples collected from 3 locations in the United States and from 2 locations in Dhaka, Bangladesh. (**B**) Network phylogeny analysis of phage genomes, compared according to shared gene content using Phamerator, as described in **Methods**, reveals 3 genomically distinct phage clusters. Colored circles indicate groups of phages according to cluster assignment, assigned by vConTACT2. (**C-E**) Annotated genome maps of representative members of each cluster (SJC01, DAC15, and DAC20). Genes are represented as colored boxes and conserved domains are inlaid yellow boxes within genes. If a gene has a conserved domain, it is annotated in black text. iVireons was used to predict structural genes as described in **Methods** and are annotated in red as predicted tail, major capsid, or general structural (tail, MCP, +S, respectively). (**F-H**) Transmission electron micrographs of SJC01, DAC15, and DAC20 show morphological differences between these representatives of the phage clusters. See **Table S1** for additional details on the isolation locations and genotypic/phenotypic characterization of these phages.

Phage genomes were annotated and compared on the basis of shared gene content (pham membership) (Cresawn et al., 2011). Phams are built and expanded when a candidate protein shares ≥32.5% identity or blastp e-value ≤1e-50 with one or more existing members of the pham. A dendrogram was built based on the presence or absence of each pham in each phage, which confirmed the three distinct genome clusters (**Figs. 1E, S1-S3**). Genome maps of representatives of each of these clusters are shown in **Figs. 1F-H.** tRNAs were detected in cluster β and γ phages (n=12-13 and n=2-3, respectively) but not in cluster α phages (**Tables S1, S2**).

While there is a high degree of intra-cluster sequence identity, there are only two phams shared between clusters: pham 150 (encoding a putative thymidylate synthase) and pham 23 (encoding collagen triple helix repeats), which are shared between all cluster β and γ representatives. Consistent with observations from previously isolated phages (Hatfull and Hendrix, 2011), the majority (roughly 80%) of phams in these *B. thetaiotaomicron*-infecting phages have no detectable conserved domains or known functions (**Tables S3-S5).**

### *Comparative analysis of* B. thetaiotaomicron *phages with existing* Bacteroides *phage isolates*

We compared these 27 *B. thetaiotaomicron-infecting* phages to 4 other previously sequenced *Bacteroides-infecting* phages (Benler et al., 2018, Ogilvie et al., 2012, Hawkins et al., 2008, Shkoporov et al., 2018) (**Fig. 2**). We noted extensive shared phams (n=53) and genome organization between the cluster β phages (DAC15 and DAC17) and CrAss001 (**Fig. 2, Fig. S4, Table S6**), reinforcing previous predictions that at least a subset of crAss-like phages prey on *Bacteroides*. A small number of phams are shared between the other isolated *B. thetaiotaomicron-infecting* phages and the previously isolated *Bacteroides-infecting* phages. Pham 22 is present in CrAss001 and the cluster γ phages. Pham 400 is present in cluster γ phages and in B124-14/B40-8, pham 887 is shared between B124-14 and the cluster α phages, pham 423 is shared between cluster α phages and B40-8, and pham 394 is shared between Hankyphage and the cluster α phages (**Table S6**). Based on this lack of relatedness, our cluster α and cluster γ phages represent the first isolates of two novel clades of *Bacteroides-infecting* phages. Furthermore, B40-8 and B124-14 are members of a separate cluster (cluster δ) and Hankyphage is a singleton with no isolated relatives (**Fig. 2**). These cluster assignments are validated by vConTACT2 (Bin Jang et al., 2019) (see **Methods**). No RefSeq phage genomes from the ProkaryoticViralRefSeq94-Merged database were grouped into clusters with these 31 isolated phages.

**Figure 2.**
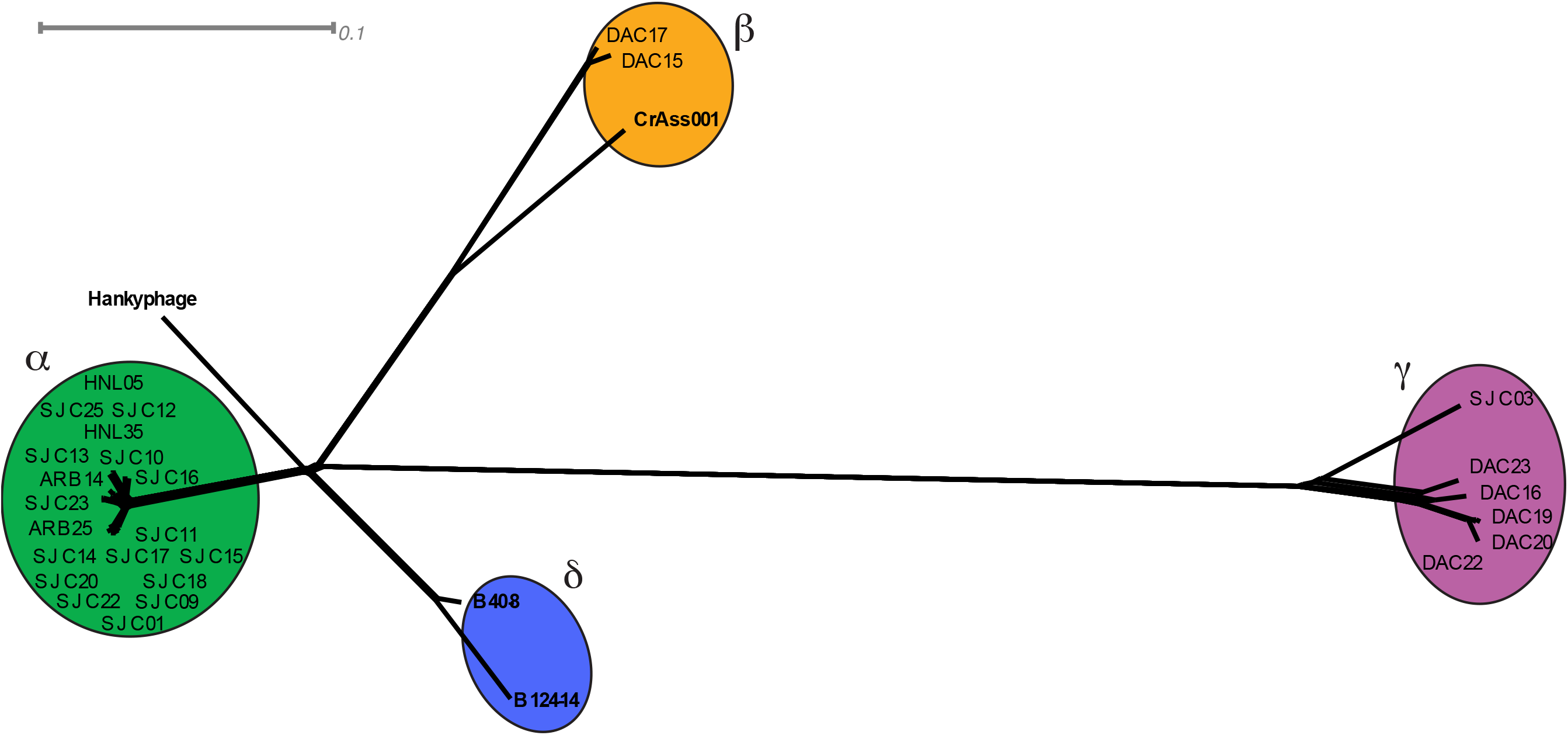
Network phylogeny of 31 *Bacteroides*-infecting phages based on gene content. The genomes of 31 *Bacteroides* infecting phages were compared according to shared gene content using Splitstree and cluster assignments were made using vConTACT2, as described in **Methods**.

### *Identification of phages related to isolated* B. thetaiotaomicron *phages in existing metagenomes*

Because the majority of phage-focused work in the gut microbiome field is based on metagenomic sequencing, we wondered if relatives of the sequenced *B. thetaiotaomicron-infecting* phage isolates could be found in existing metagenomes. To identify relatives of these phages, we used the protein search feature of SearchSRA (Torres et al., 2017, Levi et al., 2018, Towns et al., 2014, Stewart et al., 2015, Buchfink et al., 2015b, Langmead and Salzberg, 2012) to map 100,000 subsampled reads from each of the ~100,000 metagenomes in the Sequence Read Archive (SRA) onto representatives of clusters α, β, and γ (SJC01, DAC15, and DAC20, respectively). We identified 812 candidate metagenomes in the SRA where at least one of the representative phage genomes was covered by reads at an estimated real depth of >15% (given the true sequencing depth of the sample) and the percent of the genome detected was >20% for SJC01, DAC15, or DAC20 (**Fig. 3A-C**). We subsequently focused on human gut-derived metagenomes possessing sequences that are SJC01-like (>50% detected, >30x estimated coverage), DAC15-like (>40% detected, >15x estimated coverage), or DAC20-like (>20% detected) genomes for further analysis (**Table S7**). These metagenomes were downloaded from NCBI and assembled. Contigs containing significant hits (blastp e-value <1e-3) for >25% of the genes in SJC01, DAC15, or DAC20 were compared to the genomes of the isolated *Bacteroides-infecting* phages described above. See Methods for a more detailed description of this method of identifying <u>Ph</u>age <u>i</u>n <u>S</u>earc<u>h</u>SRA (PhiSh). Several PhiSh genomes were identified which are related to SJC01. These genomes include previously uncharacterized contigs from prior studies (PhiSh01 – PhiSh03, PhiSh05 – PhiSh07)(Monaco et al., 2016, He et al., 2017, Liu et al., 2016, Zheng et al., 2017, Guthrie et al., 2017) and a genome previously identified in a study examining the rapid evolution of the human gut virome (PhiSh04) (Minot et al., 2013). Serendipitously, we noticed that HSC01, a genome of a phage predicted to infect *Bacteroides caccae* (Reyes et al., 2013) is related to these cluster α isolates and cluster α-like PhiSh genomes (**Figs. 3DE; Table S8**). vConTACT2 also places all of these SJC01-like PhiSh genomes within cluster α.

**Figure 3.**
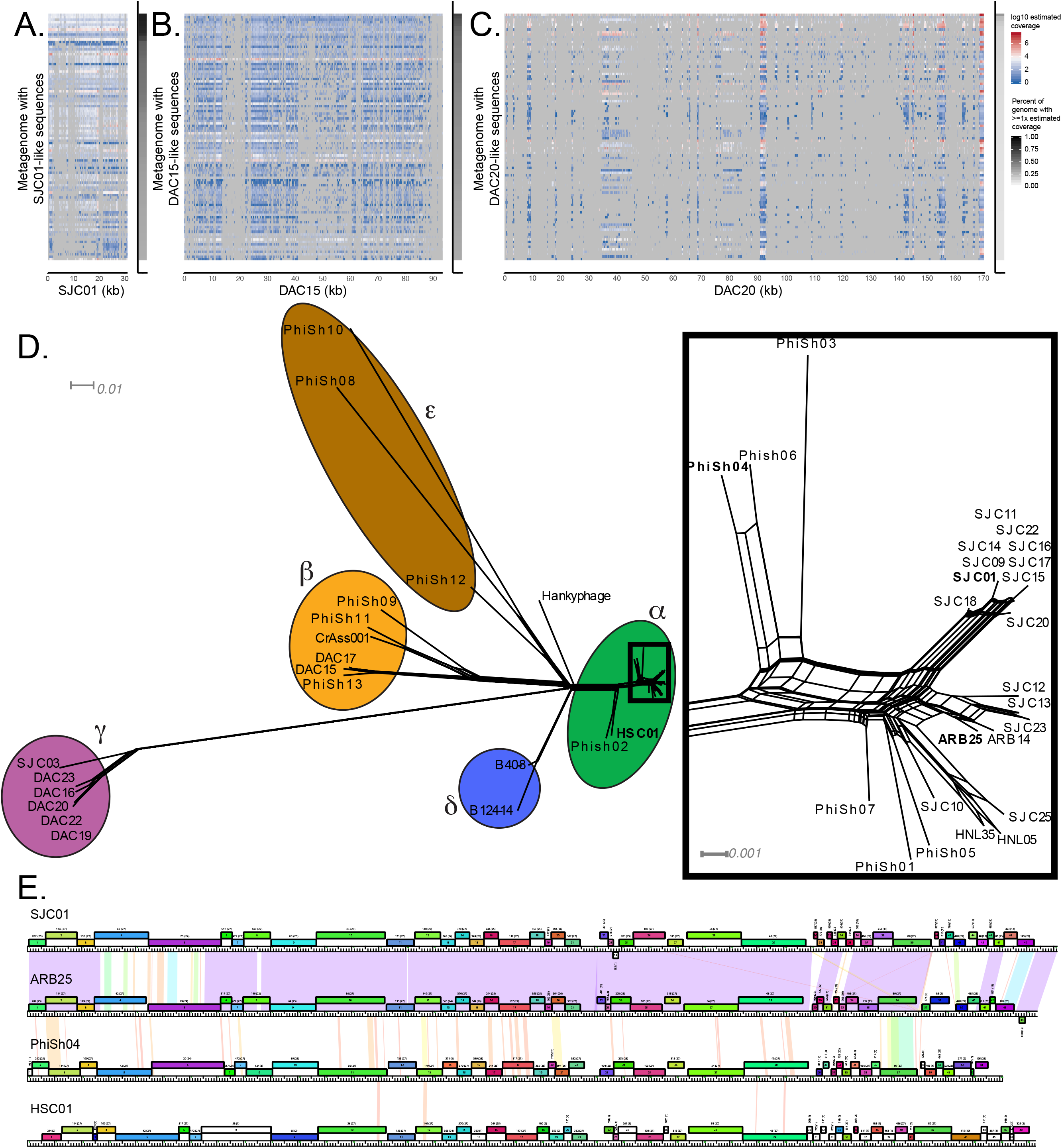
Identification of <u>Ph</u>age <u>i</u>n <u>S</u>earc<u>h</u>SRA (PhiSh) related to isolated *B. thetaiotaomicron-infecting* phages. Representatives of each genome cluster (SJC01, DAC15, DAC20) were used to query the entire NCBI SRA using SearchSRA as described in Methods. (**A-C**) Coverage depth (log10-transformed) of SJC01, DAC15, and DAC20 genomes, respectively in the 100 best hits to SJC01 identified via Search SRA (tDNA mode). The percentage of SJC01, DAC15, and DAC20 genomes detected (≥ 1 read) in each metagenome is indicated by the gray shaded column on the right of each panel. (**D**) Network phylogeny of *Bacteroides-infecting* phage genomes described in **Figure 2** and related genomes identified in publicly available metagenomes. Genomes were clustered using vConTACT2. The subset of cluster alpha phages enclosed in a rectangle is shown in greater detail at the right-hand side of the panel. Phages highlighted in panel E are in bold. (**E**) Genome maps of 4 cluster α phages (SJC01, ARB25, PhiSh04, and HSC01). The genes are color-coded according to pham membership and are numbered. Pairwise nucleotide identity is represented as shading between genomes. The color of this shading represents the degree of sequence similarity with violet being the most similar, progressing through the color spectrum to red, which is the least similar. Regions with no shading indicate no similarity with a BLASTN score of 10^-4^ or greater.

Six DAC15-like genomes were identified with this method (PhiSh08 – PhiSh13) (**Table S8**). Five of these genomes (PhiSh08 – PhiSh12) were previously identified in a study aimed at identifying crAss-like phages in human fecal metagenomes (Guerin et al., 2018) while PhiSh13 represents a novel crAss-like phage genome (He et al., 2017). Importantly, these DAC15-like PhiSh genomes are diverse (they can be classified into the previously described candidate crAss-like genera 6, 7, and 10; **Table S8**) and are differentially clustered by vConTACT2 (clusters β and ε), demonstrating that the PhiSh identification approach is capable of detecting genomes that are closely and distantly related to the PhiSh bait genome used (**Fig. 3D**).

Despite identifying a diverse collection of cluster α-like and cluster β-like PhiSh genomes, only partial γ-like PhiSh genomes were identified (**Fig. 3C**). The lack of full-length γ-like PhiSh genomes may be due to insufficient sequencing depth of the original studies or the presence of highly divergent phages which share subsets of genes with cluster γ phages.

### Identification of infection-associated phams

Our previous work demonstrated that multiple phase-variable mechanisms, including capsular polysaccharides (*cps*), modify bacteriophage susceptibility in *B. thetaiotaomicron* (Porter et al., 2019). However, phage-encoded determinants of host tropism in these phages were previously unexplored. When the *cps* specificities of these phages are compared with genome cluster membership (**Fig. 1B, Table S1**), relationships between genome cluster membership and host range become evident (**Fig. 4A**). For example, cluster γ phages tend to be most restrictive in their host range, primarily infecting *cps7, cps8*, and acapsular *B. thetaiotaomicron*. Cluster β phages are similarly restricted in their host range but are unique in their ability to efficiently infect *B. thetaiotaomicron cps3*. Some cluster α phages have promiscuous host ranges while other cluster α phages have restrictive host ranges (more similar to those of the cluster β and cluster γ phages). This variation in host range among cluster α phages prompted us to search for phams that are associated with the different infection patterns in the cluster α phages.

**Figure 4.**
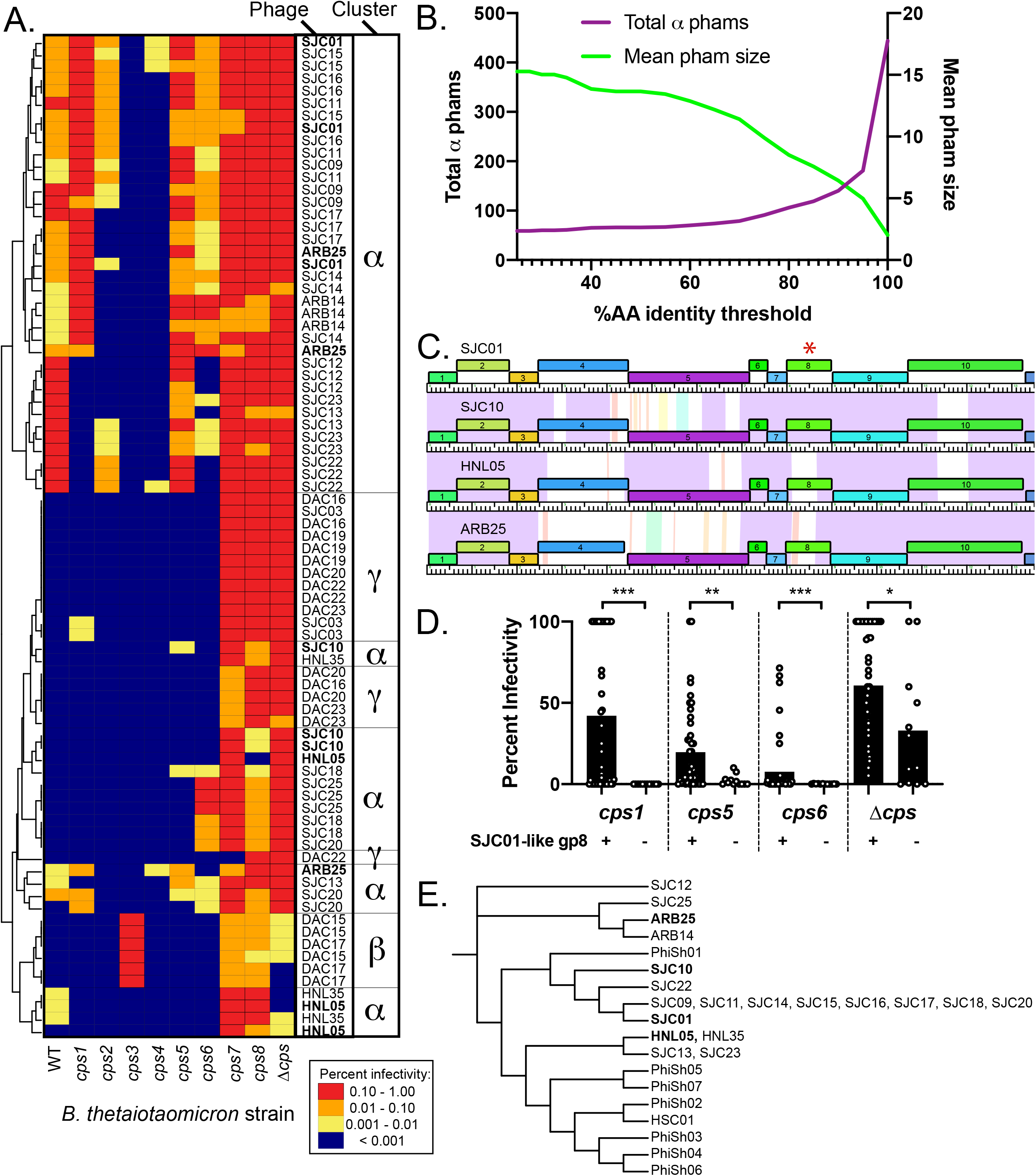
Prediction of infection-associated phams (IAPs) in *Bacteroides*-infecting phages. (**A**) Host range of *B. thetaiotaomicron* phages on strains expressing a variety of CPS (WT, wild type), a single CPS (cps1-cps8 strains) or no CPS (Δcps, acapsular). Tenfold serial dilutions of phage lysates ranging from approximately 10^6^ to 10^3^ plaque-forming units (PFU) / mL were spotted onto top agar plates containing each of the 10 bacterial strains. Plates were then incubated overnight, and plaques on each host were counted. Phage titers (PFU/ml) were calculated for each host and normalized to the titer on the “preferred host strain” for each replicate (individual replicates are shown, n=3 per phage). The phages were then clustered based on their plaquing efficiencies on the different strains (see **Methods**). Each row in the heatmap corresponds to one of three individual experimental replicates with a phage, whereas each column corresponds to one of the 10 host strains. (**B**) Changes in the total number of phams and average pham size as a function of percent amino acid identity. (**C**) Partial genome maps of 4 cluster α phages (SJC01, SJC10, HNL05, and ARB25) highlighting variation in gp4, gp5, and gp8. The genes are color coded according to pham membership at standard cutoffs and are numbered. Pairwise nucleotide identity is represented as shading between genomes. The color of this shading represents the degree of sequence similarity with violet being the most similar, progressing through the color spectrum to red, which is the least similar. Regions with no shading indicate no similarity with a BLASTN score of 10^-4^ or greater. The red asterisk highlights gp8 from these phages. Data corresponding to these 4 phages in panels A and E are in bold. (**D**) Phages containing SJC01-like gp8 were compared against phages containing the alternative allele of gp8 (85% AA identity threshold) in terms of infectivity on bacterial strains highlighted in panel A. SJC01 gp8 is associated with higher infectivity of *B. thetaiotaomicron cps1, cps5, cps6*, and Δ*cps* as assessed by Mann-Whitney U Test (p<0.05 = *, p<0.01 = **, p<0.001 = ***). (E) gp8 from cluster α isolates and the gene in the same position in cluster α genomes identified from metagenomes (PhiSh01-07, and HSC01) were aligned using ClustalW and a dendrogram of these alleles was created using The Interactive Tree of Life (See **Methods**).

We noted two major themes driving variation among the cluster α phages: variation between shared predicted structural components in these phages, such as gene products (gps) 4, 5, and 8; and mosaicism in genes at the 3’ end of the genomes, representing genes encoding small hypothetical proteins and genes encoding predicted DNA methylases. (**Fig. S1**). Therefore, we considered the possibility that allelic variation and presence/absence of phams could contribute to differences in host range among the phages. To account for each of these possibilities, we used an algorithm to identify infection-associated phams (IAPs). Specifically, we computed phams at alternative cutoffs such that membership was dictated by only by varying levels of amino acid (AA) identity between (25 and 100%; see **Methods**). As the threshold value increases, the total number of phams increases, with a concomitant decrease in mean pham membership (**Fig. 4B**), as previously observed (Cresawn et al., 2011). With the possibility that different thresholds may reveal allelic variants that correspond to infectivity, we compared these alternative pham tables with infection thresholds. This approach identified 662 total phams across all 64 infectivity/pham threshold comparisons in the 19 cluster α phages. Of these, 135 were identified as IAPs.

Among the IAPs is cluster α gp8, which is present in all cluster α phages, exhibits substantial sequence variation among these phages, and is predicted to encode a tail protein (**Figs. 1F, 4C, S1**). At the 85% AA identity cutoff, gp8 is grouped into two distinct phams and phages that have the SJC01-like variant infect *cps1, cps5, cps6*, and Δ*cps B. thetaiotaomicron* more efficiently than those that do not (**Fig. 4D**). Analysis of this IAP in the context of metagenome-derived cluster α-like phages reveals additional variation not represented in our isolates (**Fig. 4E**). Interestingly, the variants of this IAP in PhiSh02 and HSC01 contain Bacteroides-Associated Carbohydrate Binding Often N-terminal (BACON) domains (See (Reyes et al., 2013) and **Supplementary Data 1**). These combined observations suggest a role for this IAP in differential recognition of complex polysaccharides (e.g. capsular polysaccharides) across confirmed *Bacteroides-infecting* phages and related genomes.

## Discussion

In this work, we integrate phenotypic and genomic analysis of isolated phages with metagenomic analysis to highlight several opportunities for future study of gut-resident phages. In particular, though metagenome-focused studies of phages continue to generate tremendous insights into the composition and dynamics of viromes in the gut and other ecosystems, they are limited in scope due to a lack of definitive connections between predicted phages and their bacterial hosts. Several approaches have been developed to predict phage host range (Edwards et al., 2016). These approaches have been validated in part, notably for CrAss001, which was isolated on *B. intestinalis* after predictions that crAss-like phages infect members of the phylum Bacteroidetes (Dutilh et al., 2014, Yutin et al., 2018). We further validate these predictions with two more crAss-like phages, DAC15 and DAC17, which infect *B. thetaiotaomicron*. This brings the total number of published isolates of this highly abundant viral family to three (**Figs. 2, S4**). As more crAss-like phages are isolated, we anticipate that existing discrepancies relating to the roles of these phages in the gut (e.g. some crAss-like phages are associated with disease (Oude Munnink et al., 2014) while others are stably maintained in healthy individuals (Shkoporov et al., 2019)) can be disentangled with controlled experimental approaches.

Similarly, based on our genomic and metagenomic analysis of cluster α phages, we show that two previously reported phage genomes (PhiSh04 and HSC01) are related despite differences in temporal dynamics and predicted host range (**Fig. 3D**) (Reyes et al., 2013, Minot et al., 2013). This raises questions regarding what determinants encoded by phage or bacterial host are responsible for observed differences in host range and phage population dynamics. Some insights come from experiments using the cluster α phage ARB25 in gnotobiotic mice. ARB25 is stably maintained in a bi-colonization with its host for months and the phase-variable mechanisms used by *B. thetaiotaomicron* to evade ARB25 are dependent on the presence of CPS among other cell surface features (Porter et al., 2019). HSC01, unlike PhiSh04 or ARB25, does not stably co-exist with its predicted host (*B. caccae*) in gnotobiotic mice (Reyes et al., 2013), suggesting that although HSC01 is related to phages that are stably maintained, it may have distinct ecological impacts in the gut. Alternatively, it is possible that other members of the gut microbiome affect the relationship between a phage and its host bacterium.

By combining work that involves phage isolation, sequencing, and phenotypic characterization, with metagenomic analyses, we hope to reciprocally inform these studies (e.g., by adding phages and information on IAPs to publically available databases) and to provide the reagents necessary to experimentally test hypotheses using the broad toolkit available in the gut microbiome field (e.g., by probing phage-host interactions using gnotobiotics and molecular genetics). Future isolation efforts can be further optimized with high throughput approaches (e.g. robotics and automated liquid handling) or as part of educational efforts like those pioneered by the SEA-PHAGES program (Hanauer et al., 2017), which would simultaneously crowd source the effort while providing training opportunities for the next generation of microbiome scientists. Together, this integration will allow for a more comprehensive consideration of the interactions that occur between phages and their hosts at the population, individual, and molecular scales.

## Methods

### Bacterial strains and culture conditions

The bacterial strains used in this study are listed in **Table S9.** Frozen stocks of these strains were maintained in 25% glycerol at −80°C and were routinely cultured in an anaerobic chamber (Coy) under 5% H_2_, 10% CO_2_, 85% N_2_ at 37°C in *Bacteroides* Phage Recovery Medium (BPRM), as described previously (Porter et al., 2019): per 1 liter of broth, 10 g meat peptone, 10 g casein peptone, 2 g yeast extract, 5 g NaCl, 0.5 g L-cysteine monohydrate, 1.8 g glucose, and 0.12 g MgSO_4_ heptahydrate were added; after autoclaving and cooling to approximately 55 °C, 10 ml of 0.22 μm-filtered hemin solution (0.1% w/v in 0.02% NaOH), 1 ml of 0.22 μm-filtered 0.05 g/ml CaCl_2_ solution, and 25 ml of 0.22μm-filtered 1 M Na_2_CO_3_ solution were added. For BPRM agar plates, 15 g/L agar was added prior to autoclaving and hemin and Na_2_CO_3_ were added as above prior to pouring the plates. For BPRM top agar used in soft agar overlays, 3.5 g/L agar was added prior to autoclaving. Hemin, CaCl_2_, and Na_2_CO_3_ were added to the top agar as above immediately before conducting experiments. Bacterial strains were routinely struck from the freezer stocks onto BPRM agar and grown anaerobically for up to 2 days. A single colony was picked for each bacterial strain, inoculated into 5 mL BPRM, and grown anaerobically overnight to provide the starting culture for experiments.

### Bacteriophage isolation from primary wastewater effluent and sewer-adjacent pond water

The bacteriophages described in this study were isolated from primary wastewater effluent from the Ann Arbor, Michigan Wastewater Treatment Plant and from the San Jose-Santa Clara Regional Wastewater Treatment Facility, as described previously (Porter et al., 2019). For the current study, phages were isolated from primary wastewater effluent from the Sand Island Wastewater Treatment Plant (Honolulu, Hawaii) or from sewer-adjacent pond water in Dhaka, Bangladesh (**Table S1**). Water samples were centrifuged at 5,500 rcf for 10 minutes at room temperature to remove any remaining solids. The supernatant was then sequentially filtered through 0.45 μm and 0.22 μm polyvinylidene fluoride (PVDF) filters. This processed primary effluent was concentrated up to 500-fold via 100 kDa PVDF size exclusion columns.

Initial screening for plaques was done using a soft agar overlay method where 50 μL of the concentrated primary effluent was combined with 0.5 mL overnight culture and 4.5 mL BPRM top agar and poured onto a standard circular petri dish [100 mm x 15 mm]. Soft agar overlays were incubated anaerobically at 37 °C overnight. To promote a diverse collection of phages, no more than 5 plaques from the same plate were plaque purified and a diversity of plaque morphologies were selected as applicable.

Single, isolated plaques were picked into 100 μL phage buffer (prepared as an autoclaved solution of 5 ml of 1 M Tris pH 7.5, 5 ml of 1 M MgSO4, 2 g NaCl in 500 ml with ddH_2_O). Phages were plaque purified using a 96-well plate-based method, where serial dilutions were prepared in 96-well plates and 1 μL of each dilution was spotted onto a solidified top agar overlay. This procedure was repeated at least 3 times to plaque purify each phage.

High titer phage stocks were generated by flooding a soft agar overlay on a plate that yielded a “lacey” pattern of bacterial growth (near confluent lysis). Following overnight incubation of each plate, 5 ml of sterile phage buffer was added to the plate to re-suspend the phage. After at least 2 hours of incubation at room temperature, the lysate was spun at 5,500 rcf for 10 minutes to clear debris and then filter sterilized through a 0.22 μm PVDF filter. For more details on phages used in this work, see **Table S1**.

### Phage genome sequencing and assembly

DNA was extracted from high-titer phage lysates and sequencing libraries were prepared using the Ultra II FS Kit (New England Biolabs) or for ARB14 and ARB25, the TruSeq Nano DNA LT Kit (Illumina). Libraries were quantified using a BioAnalyzer (Agilent) and subsequently sequenced using 150-base single-end reads (Illumina MiSeq), or for ARB14 and ARB25, 250-base paired-end reads (Illumina MiSeq). Phage genomes were assembled using Geneious version 9.1.5 with default options after trimming reads with an error probability limit of 0.05. All genomes published here circularized during assembly. Phage genomes belonging to the same cluster were rearranged to have identical 5’ ends. Coverage for each assembly was calculated by mapping reads onto each assembled genome using bowtie2 (Langmead and Salzberg, 2012) (--very-sensitive) and then using jgi_summarize_ban_contig_depths from the MetaBAT2 tool (Kang et al., 2019) to calculate mean coverage depth.

### *Annotation and comparative analyses of* B. thetaiotaomicron *infecting phages*

Protein-coding genes and tRNAs were predicted and annotated using DNA-Master default parameters (http://cobamide2.pitt.edu/), which incorporates Genemark (Besemer and Borodovsky, 2005), Glimmer (Delcher et al., 1999), and tRNAscan-SE (Lowe and Eddy, 1997). To classify genes into related groups (phams) and identify conserved domains, Phamerator was used with default parameters (Cresawn et al., 2011). Phage genome ends and packaging strategies for cluster β phages were inferred using PhageTerm (Garneau et al., 2017) which identified clear direct terminal repeats (DTRs). PhageTerm was unable to identify DTRs or cohesive ends in the cluster α or γ phages, possibly indicating a headful packaging strategy. The large terminase proteins share significant similarity (BLASTP e-value <1e-3) with the PBSX-family of large terminases, which also use a headful packaging strategy (**Table S1**) (Anderson and Bott, 1985). To predict virion structural genes, iVireons was used with default parameters (Seguritan et al., 2012). Protein-coding genes were classified as “predicted structural genes” (e.g. general structural, tail, or capsid, annotated in **Fig. 1**) for genes with score 0.7 and above. To visualize genome-level relationships among phages, pham tables were processed with Janus (http://cobamide2.pitt.edu/) and Splitstree using default parameters. Phage genomes were clustered together using vConTACT2 and the ProkaryoticViralRefSeq94-Merged database with default parameters (Bin Jang et al., 2019). CRISPR protospacers were identified and used as the basis for host prediction of the isolated *B. thetaiotaomicron* phages and PhiSh genomes with CRISPRdb (Grissa et al., 2007) and the JGI IMG/VR Spacer Database(Paez-Espino et al., 2019) with an E-value cutoff of 1. Matches with the highest percent sequence identity are shown in **Tables S1 and S8**. Genomes of the phage isolates described in **Table S1** are uploaded to NCBI (BioProject ID PRJNA606391).

### Quantitative host range analysis

Host range analysis was carried out as previously described (Porter et al., 2019). Briefly, high titer phage stocks were prepared on their “preferred host strain,” which is the strain yielding the highest titer of phages in a pre-screen of phage host range (**Table S1**). Lysates were then diluted to approximately 10^6^ PFU/mL, were added to the wells of a 96-well plate, then further diluted to 10^5^, 10^4^, and 10^3^ PFU/mL. One microliter of each dilution was plated onto solidified top agar overlays containing wildtype *B. thetaiotaomicron*, acapsular *B. thetaiotaomicron*, or *B. thetaiotaomicron* expressing a single capsule (**Table S9**). After spots dried, plates were incubated anaerobically for 15-24 hours prior to counting plaques. Phage titers were normalized to the “preferred host strain.”

### Infection associated pham identification

We defined an infection-associated pham (IAP) as a pham that (1) was found in every phage of a given cluster (α, β, and γ; see **Fig. 1**) that infected the *B. thetaiotaomicron* isolate in question, but (2) was not found in every phage of the same cluster. Criterion (1) is a stringent threshold. For example, if 10 different phages infected a given bacterial strain, but only 9 shared a particular pham, it would fail criterion (1). Criterion (2) was included to eliminate core genes.

We employed two important thresholds when identifying IAPs. The first of these is an infection threshold - the normalized percentage of infectivity a given phage on a given isolate as described in the methods section ‘Quantitative host range analysis’. Here, a stringent threshold is 100%, which considers “infection” to be a case where the phage generates as many plaques on a given *B. thetaiotaomicron* strain as it does on its preferred host strain. A permissive threshold is 1% - here a phage would have to cause 1/100th as many plaques as it did on its preferred host. The second of these is the pham identity threshold - the percentage sequence identity that two genes must share to be counted as in the same pham. This clustering is described in methods section ‘Annotation and comparative analyses of *B. thetaiotaomicron* infecting phages.’ Here, a stringent clustering threshold is 100%, where genes sharing 100% sequence identity are grouped in the same pham. A permissive threshold would be 1%. The lower this threshold, the more disparate the sequences that are grouped together.

We computed our IAP identification algorithm using as thresholds each member of the product set of [1%, 5%, 10%, 50%] X [25%, 27.5%, 30%, 35%, 40%, 45%, 50%, 60%, 70%, 75%, 80%, 85%, 90%, 95%, 100%] (infection threshold and pham identity threshold, respectively). Code and data are available as **Supplementary Data 2**, which provides a simple python script and the accompanying data allowing exact reproduction of the method.

Comparisons of cluster α gp8 and homologs from metagenome-derived cluster α genomes (PhiSh01-PhiSh07, HSC01) were conducted using Clustal Omega (Sievers et al., 2011) and visualized using The Interactive Tree of Life (Letunic and Bork, 2019) with default parameters.

### Transmission Electron Microscopy

High titer phage lysates of representatives from each genome cluster (SJC01, DAC15, DAC20) were precipitated overnight at 4°C with gentle rocking in a solution of 1M NaCl and 10% w/v PEG8000. Phages were then precipitated via centrifugation (5500xg for 10 minutes at 4°C). Six milliliters of phage buffer was added to the pellet and broken with gentle agitation and swirling and the mixture was incubated overnight at 4°C with gentle rocking. The following day, the sample was centrifuged at 5500xg for 10 minutes at 4°C. CsCl was slowly added to the supernatant and gently dissolved via gentle swirling (final concentration 75% w/v solution). Samples were centrifuged at 26,000 RPM for 24 hours at 5°C. Phage bands were extracted and stored at 4°C.

CsCl-banded lysates were applied directly to glow discharged Carbon Type-B 200 mesh copper grids. Samples were allowed to adsorb to the grids for 3 minutes and were subsequently washed with 2 drops of ultrapure water. Three drops of uranyl acetate (1% w/v in water) were applied to the grid and the third drop was maintained on the grid for 1 minute. Filter paper was used to remove the majority of the uranyl acetate and allowed to dry at room temperature. Samples were then viewed at 120 kV on a JEOL JEM-1400 transmission electron microscope and images were collected using a Gatan Orius digital camera.

### *Comparative genomic analyses between isolated* B. thetaiotaomicron *infecting phages, other isolated* Bacteroides*-infecting phages, and PhiSh genomes*

Genomes of representatives of each genome cluster (SJC01, DAC15, DAC20) were queried against the entire SRA using SearchSRA (Torres et al., 2017, Levi et al., 2018, Towns et al., 2014, Stewart et al., 2015, Buchfink et al., 2015b, Langmead and Salzberg, 2012). To determine whether these genome clusters are found in human gut metagenomes, one representative from each cluster (SJC01, DAC15, DAC20) was queried using SearchSRA using the “protein search” option. SearchSRA uses DIAMOND blastx to query 100,000 reads from each of ~100,000 metagenomes publicly available in NCBI SRA against a single query amino acid sequence. The input data for each representative phage genome consisted of a single amino acid sequence consisting of every translated gene in order of appearance in the genome, separated by “XXX”. This input format was required when the analysis was conducted (July 24, 2019).

Data were retrieved from SearchSRA in the typical BLAST M8 format (one file per NCBI metagenome aligned to the reference phage) and parsed into BED format. BEDTools (Quinlan, 2014) coverage was used to calculate the coverage depth of each base pair along the genome. These tables were read into R 3.6.2. For each sequence run (SRR) that had ≥1 read aligning to a query amino acid sequence, SRAdb (Zhu et al., 2013) was used to get the associated sample accession number (SRS) and other related sample metadata. Coverage data from sequencing runs belonging to the same sample were combined, and then average coverage depth and detection (% of bases with ≥ 1x coverage) was calculated for each metagenome sample mapped.

For each metagenome sample mapped where the number of reads sequenced was >10000, the estimated true coverage depth of the reference phage in that metagenome sample was calculated as # spots sequenced*SearchSRA average coverage / 100000. To determine whether to assemble a given metagenome and search for a relative of a given representative phage, we filtered the list of metagenome samples based on whether the estimated real coverage was >15% and the percent of the genome detected was >20%. This list was filtered further by selecting only human gut metagenomes and by selecting samples where coverage and detection were the highest (**Table S7**).

Metagenomes were downloaded from NCBI SRA using parallel-fastq-dump 0.6.5 (https://github.com/rvalieris/parallel-fastq-dump). For each metagenome assembled, reads were trimmed using BBDuk (https://sourceforge.net/projects/bbmap/) 38.69 (parameters ref=adapters,phix threads=$(($coreNum - 2)) ktrim=r k=23 mink=11 hdist=2 tpe tbo qtrim=rl trimq=20 minlen=55) and assembled using MEGAHIT v1.2.9 (--mem-flag 2 –k-list 21,29,39,49,59,69,79,89,99) for all samples, or –k-list 21,29,39,49,59,69,79,89,99,109,119,129,139,149 if read length was >=2×250bp.

To identify contigs in the metagenome assemblies that might be putative relatives of the representative phages, we used DIAMOND 0.9.24(Buchfink et al., 2015a) to build a blastx database containing all individual amino acid sequences from all three representative genomes. DIAMOND blastx queries consisted all contigs from a single metagenome assembly. Individual contigs containing significant (e <= 0.001) hits for >25% of the genes from a given representative phage genome were reoriented to align the 5’ ends with isolated phage genomes and then included in subsequent Phamerator analysis (**Table S8**). See **Supplementary Data 1** for Genbank and fasta files of the PhiSh genomes. A tutorial for performing this analysis can be found as **Supplementary Data 3**.

## Supplementary Data

**Supplementary Data 1-3** are accessible at https://drive.google.com/open?id=100fpin0lDy-6iGDDe17-OEzs0yrNHlBu.

## Author contributions

AJH, BDM, NTP, WVT, DAR, and RAG performed experiments/computational analyses, and analyzed the data. AJH, BDM, NTP, and WVT prepared the display items. EJN, ECM, and JLS provided key insights, tools, and reagents. AJH wrote the paper. All authors edited the manuscript prior to submission.

## Acknowledgements

We thank Jackson Gardner for assistance with host range analyses; Gayatri Vithanage, Lyle Shizumura, Greig Steward, and Ned Ruby for logistical assistance in phage isolation from Sand Island Wastewater Treatment Plant; and John Perrino for transmission electron microscopy expertise. This work was funded by NIH grants (GM099513 and DK096023 to ECM; DP5OD019893 to EJN, DK085025 and AT00989203 to JLS), an NIH postdoctoral NRSA (5T32AI007328 to AJH), a Stanford University School of Medicine Dean’s Postdoctoral Fellowship (AJH), the NIH Cellular Biotechnology Training Program (T32GM008353 to NTP), by a NCCR ARRA Award (1S10RR026780-01 to Stanford University Cell Sciences Imaging Facility), and by a National Science Foundation Graduate Research Fellowship (DGE-114747 to BDM). JLS is a Chan Zuckerberg Biohub Investigator.

---

**<u>Figure S1.</u> Genome maps of *B. thetaiotaomicron*-infecting cluster α phages.** The genes are color-coded according to pham membership and are numbered. Pairwise nucleotide identity is represented as shading between genomes. The color of this shading represents the degree of sequence similarity with violet being the most similar, progressing through the color spectrum to red, which is the least similar. Regions with no shading indicate no similarity with a BLASTN score of 10^-4^ or greater.

**<u>Figure S2.</u> Genome maps of *B. thetaiotaomicron*-infecting cluster β phages.** The genes are color-coded according to pham membership and are numbered. Pairwise nucleotide identity is represented as shading between genomes. The color of this shading represents the degree of sequence similarity with violet being the most similar, progressing through the color spectrum to red, which is the least similar. Regions with no shading indicate no similarity with a BLASTN score of 10^-4^ or greater.

**<u>Figure S3.</u> Genome maps of *B. thetaiotaomicron*-infecting cluster γ phages.** The genes are color-coded according to pham membership and are numbered. Pairwise nucleotide identity is represented as shading between genomes. The color of this shading represents the degree of sequence similarity with violet being the most similar, progressing through the color spectrum to red, which is the least similar. Regions with no shading indicate no similarity with a BLASTN score of 10^-4^ or greater.

**<u>Figure S4.</u> Genome maps of DAC15, DAC17, and CrAss001.** The genes are color coded according to pham membership and are numbered. Pairwise nucleotide identity is represented as shading between genomes. The color of this shading represents the degree of sequence similarity with violet being the most similar, progressing through the color spectrum to red, which is the least similar. Regions with no shading indicate no similarity with a BLASTN score of 10^-4^ or greater.

## References

Anderson, L. M. & Bott, K. F. 1985. DNA packaging by the *Bacillus subtilis* defective bacteriophage PBSX. J Virol, 54, 773–80.

Barr, J. J., Auro, R., Furlan, M., Whiteson, K. L., Erb, M. L., Pogliano, J., Stotland, A., Wolkowicz, R., Cutting, A. S., Doran, K. S., Salamon, P., Youle, M. & Rohwer, F. 2013. Bacteriophage adhering to mucus provide a non-host-derived immunity. Proc Natl Acad Sci U S A, 110, 10771–6.

Benler, S., Cobian-Guemes, A. G., Mcnair, K., Hung, S. H., Levi, K., Edwards, R. & Rohwer, F. 2018. A diversity-generating retroelement encoded by a globally ubiquitous Bacteroides phage. Microbiome, 6, 191.

Besemer, J. & Borodovsky, M. 2005. GeneMark: web software for gene finding in prokaryotes, eukaryotes and viruses. Nucleic Acids Res, 33, W451–4.

Bin Jang, H., Bolduc, B., Zablocki, O., Kuhn, J. H., Roux, S., Adriaenssens, E. M., Brister, J. R., Kropinski, A. M., Krupovic, M., Lavigne, R., Turner, D. & Sullivan, M. B. 2019. Taxonomic assignment of uncultivated prokaryotic virus genomes is enabled by gene-sharing networks. Nat Biotechnol, 37, 632–639.

Brussow, H. & Hendrix, R. W. 2002. Phage genomics: small is beautiful. Cell, 108, 13–6.

Buchfink, B., Xie, C. & Huson, D. H. 2015a. Fast and sensitive protein alignment using DIAMOND. Nat Methods, 12, 59–60.

Buchfink, B., Xie, C. & Huson, D. H. 2015b. Fast and sensitive protein alignment using DIAMOND. Nat Methods.

Cresawn, S. G., Bogel, M., Day, N., Jacobs-Sera, D., Hendrix, R. W. & Hatfull, G. F. 2011. Phamerator: a bioinformatic tool for comparative bacteriophage genomics. BMC Bioinformatics, 12, 395.

Delcher, A. L., Harmon, D., Kasif, S., White, O. & Salzberg, S. L. 1999. Improved microbial gene identification with GLIMMER. Nucleic Acids Res, 27, 4636–41.

Duerkop, B. A., Kleiner, M., Paez-Espino, D., Zhu, W., Bushnell, B., Hassell, B., Winter, S. E., Kyrpides, N. C. & Hooper, L. V. 2018. Murine colitis reveals a disease-associated bacteriophage community. Nat Microbiol.

Dutilh, B. E., Cassman, N., Mcnair, K., Sanchez, S. E., Silva, G. G., Boling, L., Barr, J. J., Speth, D. R., Seguritan, V., Aziz, R. K., Felts, B., Dinsdale, E. A., Mokili, J. L. & Edwards, R. A. 2014. A highly abundant bacteriophage discovered in the unknown sequences of human faecal metagenomes. Nat Commun, 5, 4498.

Edwards, R. A., Mcnair, K., Faust, K., Raes, J. & Dutilh, B. E. 2016. Computational approaches to predict bacteriophage-host relationships. FEMS Microbiol Rev, 40, 258–72.

Garneau, J. R., Depardieu, F., Fortier, L. C., Bikard, D. & Monot, M. 2017. PhageTerm: a tool for fast and accurate determination of phage termini and packaging mechanism using next-generation sequencing data. Sci Rep, 7, 8292.

Grazziotin, A. L., Koonin, E. V. & Kristensen, D. M. 2017. Prokaryotic Virus Orthologous Groups (pVOGs): a resource for comparative genomics and protein family annotation. Nucleic Acids Res, 45, D491–d498.

Gregory, A. C., Zablocki, O., Howell, A., Bolduc, B. & Sullivan, M. B. 2019. The human gut virome database. bioRxiv.

Grissa, I., Vergnaud, G. & Pourcel, C. 2007. The CRISPRdb database and tools to display CRISPRs and to generate dictionaries of spacers and repeats. BMC Bioinformatics, 8, 172.

Guerin, E., Shkoporov, A., Stockdale, S. R., Clooney, A. G., Ryan, F. J., Sutton, T. D. S., Draper, L. A., Gonzalez-Tortuero, E., Ross, R. P. & Hill, C. 2018. Biology and Taxonomy of crAss-like Bacteriophages, the Most Abundant Virus in the Human Gut. Cell Host Microbe, 24, 653–664. e6.

Guthrie, L., Gupta, S., Daily, J. & Kelly, L. 2017. Human microbiome signatures of differential colorectal cancer drug metabolism. NPJ Biofilms Microbiomes, 3, 27.

Hanauer, D. I., Graham, M. J., Betancur, L., Bobrownicki, A., Cresawn, S. G., Garlena, R. A., Jacobs-Sera, D., Kaufmann, N., Pope, W. H., Russell, D. A., Jacobs, W. R., JR., Sivanathan, V., Asai, D. J. & Hatfull, G. F. 2017. An inclusive Research Education Community (iREC): Impact of the SEA-PHAGES program on research outcomes and student learning. Proc Natl Acad Sci U S A, 114, 13531–13536.

Hatfull, G. F. & Hendrix, R. W. 2011. Bacteriophages and their genomes. Curr Opin Virol, 1, 298–303.

Hawkins, S. A., Layton, A. C., Ripp, S., Williams, D. & Sayler, G. S. 2008. Genome sequence of the *Bacteroides fragilis* phage ATCC 51477-B1. Virol J, 5, 97.

He, Q., Gao, Y., Jie, Z., Yu, X., Laursen, J. M., Xiao, L., Li, Y., Li, L., Zhang, F., Feng, Q., Li, X., Yu, J., Liu, C., Lan, P., Yan, T., Liu, X., Xu, X., Yang, H., Wang, J., Madsen, L., Brix, S., Wang, J., Kristiansen, K. & Jia, H. 2017. Two distinct metacommunities characterize the gut microbiota in Crohn’s disease patients. Gigascience, 6, 1–11.

Kang, D. D., Li, F., Kirton, E., Thomas, A., Egan, R., An, H. & Wang, Z. 2019. MetaBAT 2: an adaptive binning algorithm for robust and efficient genome reconstruction from metagenome assemblies. PeerJ, 7, e7359.

Langmead, B. & Salzberg, S. L. 2012. Fast gapped-read alignment with Bowtie 2. Nat Methods

Letunic, I. & Bork, P. 2019. Interactive Tree Of Life (iTOL) v4: recent updates and new developments. Nucleic Acids Res, 47, W256–w259.

Levi, K., Rynge, M., Eroma, A. & Edwards, R. A. 2018. Searching the Sequence Read Archive using Jetstream and Wrangler. Proceedings of the Practice and Experience on Advanced Research Computing.

Liu, W., Zhang, J., Wu, C., Cai, S., Huang, W., Chen, J., Xi, X., Liang, Z., Hou, Q., Zhou, B., Qin, N. & Zhang, H. 2016. Unique Features of Ethnic Mongolian Gut Microbiome revealed by metagenomic analysis. Sci Rep, 6, 34826.

Lowe, T. M. & Eddy, S. R. 1997. tRNAscan-SE: a program for improved detection of transfer RNA genes in genomic sequence. Nucleic Acids Res, 25, 955–64.

Manrique, P., Bolduc, B., Walk, S. T., Van Der Oost, J., De Vos, W. M. & Young, M. J. 2016. Healthy human gut phageome. Proc Natl Acad Sci U S A, 113, 10400–5.

Minot, S., Bryson, A., Chehoud, C., Wu, G. D., Lewis, J. D. & Bushman, F. D. 2013. Rapid evolution of the human gut virome. Proc Natl Acad Sci U S A, 110, 12450–5.

Monaco, C. L., Gootenberg, D. B., Zhao, G., Handley, S. A., Ghebremichael, M. S., Lim, E. S., Lankowski, A., Baldridge, M. T., Wilen, C. B., Flagg, M., Norman, J. M., Keller, B. C., Luevano, J. M., Wang, D., Boum, Y., Martin, J. N., Hunt, P. W., Bangsberg, D. R., Siedner, M. J., Kwon, D. S. & Virgin, H. W. 2016. Altered Virome and Bacterial Microbiome in Human Immunodeficiency Virus-Associated Acquired Immunodeficiency Syndrome. Cell Host Microbe, 19, 311–22.

Ogilvie, L. A., Caplin, J., Dedi, C., Diston, D., Cheek, E., Bowler, L., Taylor, H., Ebdon, J. & Jones, B. V. 2012. Comparative (meta)genomic analysis and ecological profiling of human gut-specific bacteriophage phiB124-14. PLoS One, 7, e35053.

Oude Munnink, B. B., Canuti, M., Deijs, M., De Vries, M., Jebbink, M. F., Rebers, S., Molenkamp, R., Van Hemert, F. J., Chung, K., Cotten, M., Snijders, F., Sol, C. J. & Van Der Hoek, L. 2014. Unexplained diarrhoea in HIV-1 infected individuals. BMC Infect Dis, 14, 22.

Paez-Espino, D., Roux, S., Chen, I. A., Palaniappan, K., Ratner, A., Chu, K., Huntemann, M., Reddy, T. B. K., Pons, J. C., Llabres, M., Eloe-Fadrosh, E. A., Ivanova, N. N. & Kyrpides, N. C. 2019. IMG/VR v.2.0: an integrated data management and analysis system for cultivated and environmental viral genomes. Nucleic Acids Res, 47, D678–d686.

Porter, N. T., Hryckowian, A. J., Merrill, B. D., Gardner, J. O., Singh, S., Sonnenburg, J. L. & Martens, E. C. 2019. Multiple phase-variable mechanisms, including capsular polysaccharides, modify bacteriophage susceptibility in *Bacteroides thetaiotaomicron*. bioRxiv.

Quinlan, A. R. 2014. BEDTools: The Swiss-Army Tool for Genome Feature Analysis. Curr Protoc Bioinformatics, 47, 11.12.1–34.

Ren, J., Ahlgren, N. A., Lu, Y. Y., Fuhrman, J. A. & Sun, F. 2017. VirFinder: a novel k-mer based tool for identifying viral sequences from assembled metagenomic data. Microbiome, 5, 69.

Reyes, A., Wu, M., Mcnulty, N. P., Rohwer, F. L. & Gordon, J. I. 2013. Gnotobiotic mouse model of phage-bacterial host dynamics in the human gut. Proc Natl Acad Sci U S A, 110, 20236–41.

Roux, S., Enault, F., Hurwitz, B. L. & Sullivan, M. B. 2015. VirSorter: mining viral signal from microbial genomic data. PeerJ, 3, e985.

Seguritan, V., Alves, N., JR., Arnoult, M., Raymond, A., Lorimer, D., Burgin, A. B., JR., Salamon, P. & Segall, A. M. 2012. Artificial neural networks trained to detect viral and phage structural proteins. PLoS Comput Biol, 8, e1002657.

Shkoporov, A. N., Clooney, A. G., Sutton, T. D. S., Ryan, F. J., Daly, K. M., Nolan, J. A., Mcdonnell, S. A., Khokhlova, E. V., Draper, L. A., Forde, A., Guerin, E., Velayudhan, V., Ross, R. P. & Hill, C. 2019. The Human Gut Virome Is Highly Diverse, Stable, and Individual Specific. Cell Host Microbe, 26, 527–541. e5.

Shkoporov, A. N., Khokhlova, E. V., Fitzgerald, C. B., Stockdale, S. R., Draper, L. A., Ross, R. P. & Hill, C. 2018. PhiCrAss001 represents the most abundant bacteriophage family in the human gut and infects Bacteroides intestinalis. Nat Commun, 9, 4781.

Sievers, F., Wilm, A., Dineen, D., Gibson, T. J., Karplus, K., Li, W., Lopez, R., Mcwilliam, H., Remmert, M., Soding, J., Thompson, J. D. & Higgins, D. G. 2011. Fast, scalable generation of high-quality protein multiple sequence alignments using Clustal Omega. Mol Syst Biol, 7, 539.

Stewart, C. A., Cockerill, T. M., Foster, I., Hancock, D., Merchant, N., Skidmore, E., Stanzione, D., Taylor, J., Tuecke, S., Turner, G., Vaughn, M. & Gaffney, N. I. 2015. Jetstream: a self-provisioned, scalable science and engineering cloud environment. Proceedings of the 2015 XSEDE Conference: Scientific Advancements Enabled by Enhanced Cyberinfrastructure.

Torres, P. J., Edwards, R. A. & Mcnair, K. A. 2017. {PARTIE}: a partition engine to separate metagenomic and amplicon projects in the Sequence Read Archive. Bioinformatics.

Towns, J., Cockerill, T., Dahan, M., Foster, I., Gaither, K., Grimshaw, A., Hazlewood, V., Lathrop, S., Lifka, D., Peterson, G. D., Roskies, R., Scott, J. R. & Wilkins-Diehr, N. 2014. XSEDE: Accelerating Scientific Discovery. Computing in Science Engineering.

Yutin, N., Makarova, K. S., Gussow, A. B., Krupovic, M., Segall, A., Edwards, R. A. & Koonin, E. V. 2018. Discovery of an expansive bacteriophage family that includes the most abundant viruses from the human gut. Nat Microbiol, 3, 38–46.

Zheng, S., Shao, S., Qiao, Z., Chen, X., Piao, C., Yu, Y., Gao, F., Zhang, J. & Du, J. 2017. Clinical Parameters and Gut Microbiome Changes Before and After Surgery in Thoracic Aortic Dissection in Patients with Gastrointestinal Complications. Sci Rep, 7, 15228.

Zhu, Y., Stephens, R. M., Meltzer, P. S. & Davis, S. R. 2013. SRAdb: query and use public next-generation sequencing data from within R. BMC Bioinformatics, 14, 19.

