## Supplementary figures and images for "*Bacteroides thetaiotaomicron*-infecting bacteriophage isolates inform sequence-based host range predictions"

### Supplementary Figure 1

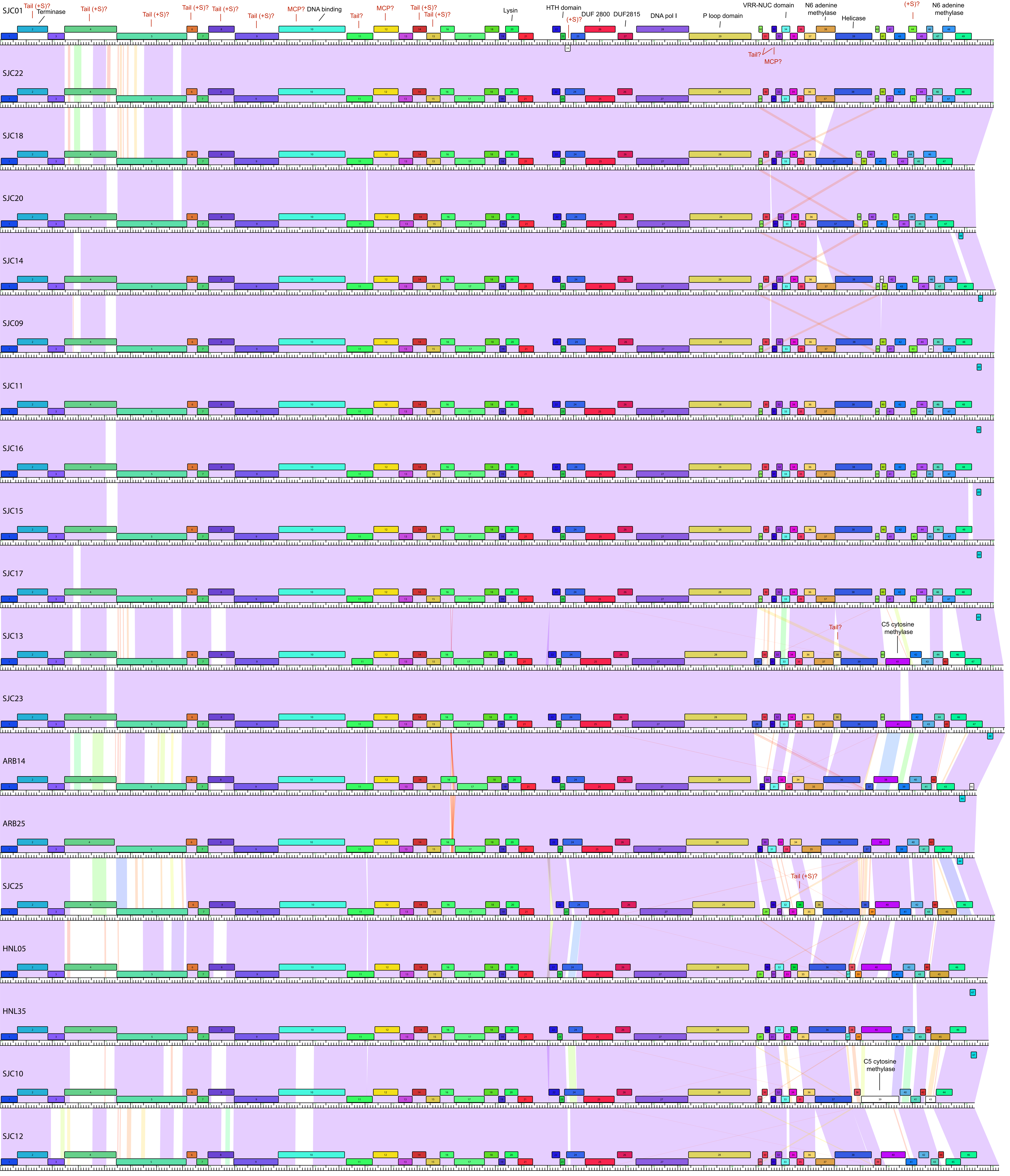

### Supplementary Figure 2

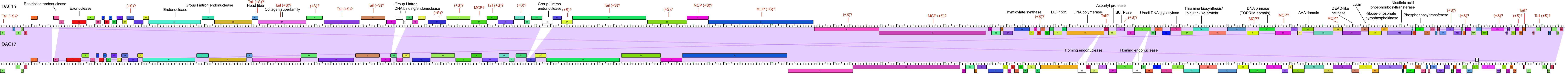

### Supplementary Figure 3

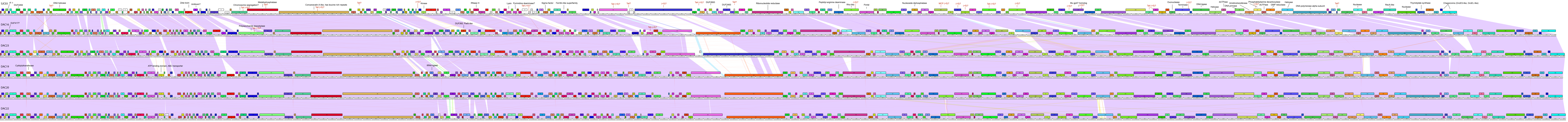

### Supplementary Figure 4

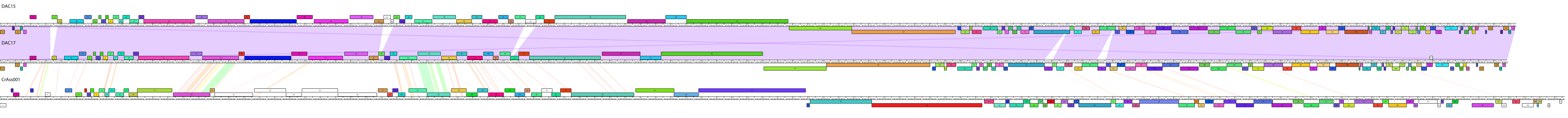
